# Current challenges in GWAS integration and fine-mapping for variant interpretation

**DOI:** 10.64898/2026.07.04.736511

**Authors:** Omar Y. Ahmed, Neha Saravanan, Anne B. Rovsing, Danny Simpson, Archit Devarajan, Sophia Gunn, Tarjinder Singh, Tuuli Lappalainen, Neville E. Sanjana

## Abstract

Over the past two decades, genome-wide association studies (GWAS) have identified thousands of trait- and disease-associated loci. However, the mechanistic understanding of these loci remains incomplete, which limits our ability to understand gene regulation and cellular programs underlying complex traits, predict disease risk, and develop therapeutics targeted to root causes. Here, we describe the current challenges for using GWAS to prioritize variants for functional follow-up experiments. These challenges span multiple domains, including limitations in data sharing and harmonization, limitations of statistical and functional fine-mapping, and the ambiguity in the added value of emerging deep learning frameworks for variant effect prediction as a complementary approach alongside traditional statistical genetics methods. We analyze these variant prioritization methods and suggest a multi-modal approach for resolving GWAS loci to a focused set of high-confidence variants for functional exploration. Fully realizing the potential of GWAS will require harmonized summary statistics and broader sharing of in-sample linkage disequilibrium (LD) data to enable robust and scalable causal variant prioritization.

## Main

Genome-wide association studies (GWAS) have identified over 625,000 unique variant-trait links across ∼15,000 phenotypes, providing a potential causal link between molecular variation and disease^1,2^. Clinical trials for drug targets with genetic evidence are three times more likely to experience clinical success, underscoring the importance of fully utilizing GWAS resources^3^. For example, the first approved gene editing therapy (for sickle-cell anemia and beta thalassemia) targets a GWAS locus near *BCL11A* associated with persistent fetal hemoglobin (HbF) in adults^4^. Mechanistic studies identified an erythroid-specific enhancer in this locus that when disrupted with CRISPR gene editing increases HbF expression and mitigates clinical manifestations of the disease^5–7^. However, molecular mechanisms have been identified for only a small fraction of GWAS loci, and even fewer have led to new treatments. The analytical and experimental strategies to best utilize the rich GWAS resources are still under development.

In GWAS, most trait-associated variants reside in the non-coding regions of the genome, making it difficult to assess their underlying mechanism^2,8^. In contrast, the functional impact of coding variants can be determined with relative ease as numerous tools have been developed for this task^9–11^. Another fundamental challenge is that GWAS identifies loci where linkage disequilibrium (LD) often makes it difficult to distinguish which variants are driving the association, and which variants are simply correlated with the causal variant. In many loci, there may not be a single causal variant but instead multiple functional variants in LD, as suggested by recent computational analyses and experimental enhancer validation via reporter assays^12–14^.

Functional analysis of GWAS loci has largely focused on identifying the target genes in the locus, with less emphasis on the causal variants themselves. However, identifying causal variants improves the portability of polygenic risk scores across different ancestries^15,16^, enhances the detection of natural selection in human populations^17,18^, helps identify key nodes within complex gene regulatory networks^19^, and pinpoints transcription factors whose altered binding drives disease risk^20^. In addition, these variants give us the molecular blueprints to move beyond gene-targeting to a more precise intervention, as exemplified by the gene editing therapy targeting an erythroid-specific enhancer: This can improve safety by altering gene expression only in a specific cell type (e.g. red blood cells and their progenitors)^5^. Importantly, the delineation of molecular mechanisms and target genes of causal noncoding GWAS variants typically requires experimental or *in silico* analysis of putative causal variants and their positions in the genome, which need to be first inferred from GWAS data. Thus, challenges in causal variant inference create a barrier for experimental follow-up of GWAS loci.

The importance of causal variant discovery, the growing number of large-scale GWAS, and increasing availability of experimental tools for functional characterization make it essential to have robust, scalable and accurate approaches to move from GWAS to causal variants (**Fig. 1a**). Here, we document the challenges in this process, focusing on the obstacles that it poses in using genetic association studies to guide downstream functional experiments. First, we detail issues and difficulties with current GWAS data sharing practices as well as simulate realistic fine-mapping conditions to highlight the consequences of poor or unmatched LD reference data. Furthermore, we examine the impact of LD mismatch on statistical fine-mapping to nominate putative causal variants, and explore how combining GWAS with gene expression data, functional annotations, and multi-study colocalization aids causal variant identification. Increasingly, more sophisticated experimental designs seek to integrate multiple GWAS studies, but leveraging their complementary strengths often comes with analytical challenges. Lastly, we apply six deep learning models to CRISPR perturbations of fine-mapped GWAS variants (blood traits from UK Biobank and the Blood Cell Consortium) to evaluate how well these frameworks predict causal variants (**Fig. 1b**). Resolving the challenges associated with inference of likely causal variants is vital for designing experiments with high signal-to-noise and that are feasible and cost-efficient, in terms of perturbations/variants tested and cells required (**Fig. 1c**). Altogether, our results show that further investments to tackle the challenges in the post-GWAS analytical pipeline will improve their downstream value for biological discovery.

**Fig. 1.**
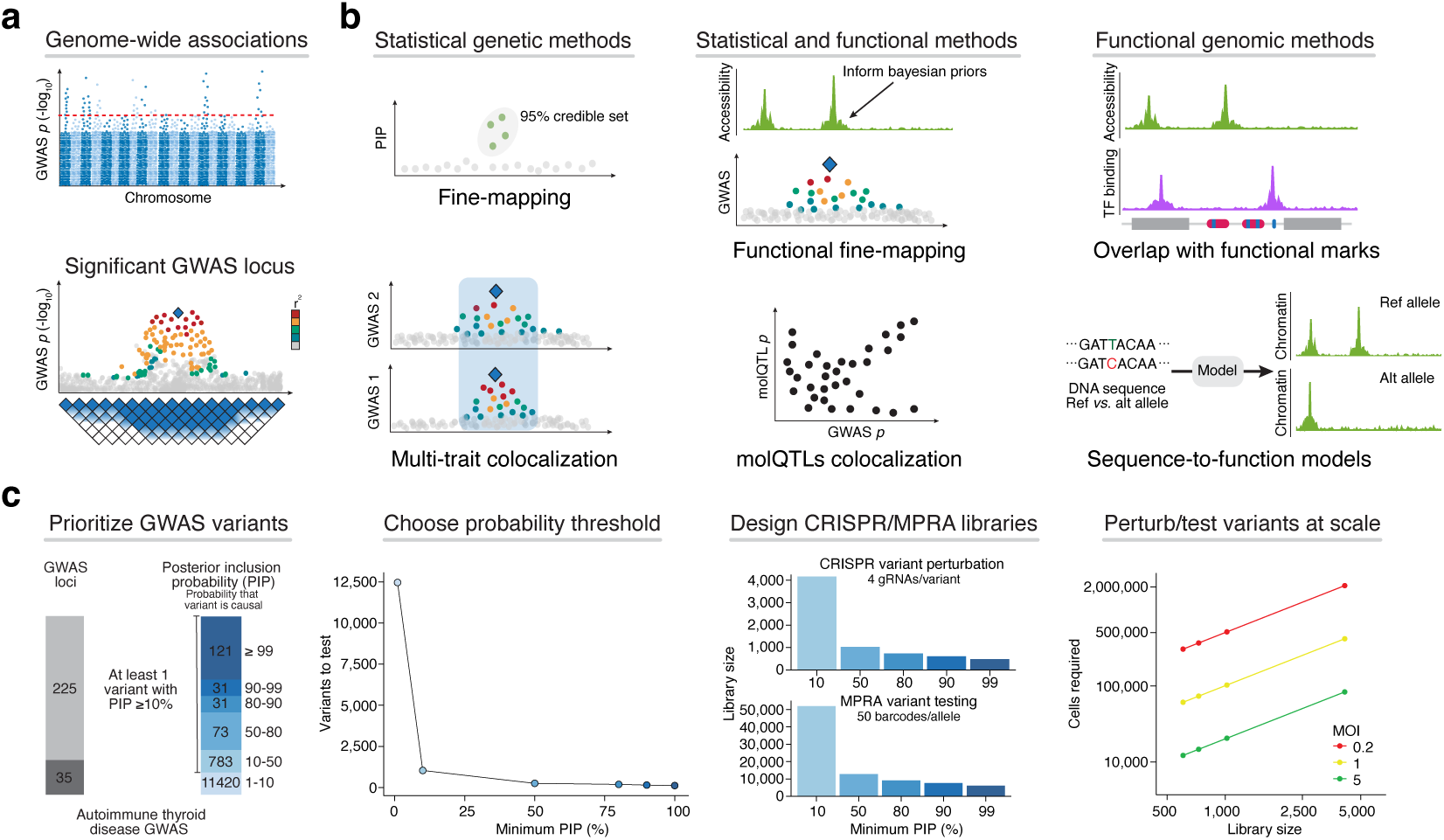
The post-GWAS discovery pipeline involves a wide array of *in-silico* and experimental approaches. **a,** Genome wide association studies (GWAS) identify loci in the genome where variants strongly associate with the phenotype. Tight linkage disequilibrium blocks complicate the identification of the causal variant. **b,** Numerous *in-silico* variant prioritization methods exist to nominate putative causal variants from GWAS data, utilizing statistical genetics and functional genomics data. **c,** Variant prioritization methods, such as fine-mapping, guide the post-GWAS discovery pipeline by ranking variants using confidence scores like posterior inclusion probability. The schematic visualizes the process of selecting variants for downstream validation experiments; we fine-mapped 260 significant GWAS loci from a recent autoimmune thyroid disease GWAS and used the results as an example^76^. Downstream validation experiments (e.g. CRISPR/MPRAs) can become cost-prohibitive with low signal-to-noise ratio as target sizes grow, therefore *in-silico* methods are crucial for trimming down the target variant set. The estimate of cells needed depends on the multiplicity of infection (MOI), which is the average number of CRISPR perturbations or MPRA variants tested per cell.

### Limitations in GWAS data sharing practices and availability

The large number of GWAS has made it valuable to compare and integrate results between studies, enabled by commendable community standards for sharing genome-wide summary statistics^1^. However, integrating data can be challenging for several reasons, including different genotyping technologies and variants measured, different genome builds, different LD structures, and different allele frequency thresholds. GWAS depend on two primary inputs: the genotyping technology and the imputation reference panel, which together determine the coverage of SNPs^21,22^. These inputs can vary widely across studies, and may even be customized for specific disease groups (e.g., ImmunoChip which is tailored toward autoimmune and inflammatory diseases^23^). Further, a single GWAS can include several different genotyping technologies: When examining 123 autoimmunity-related GWAS (covering 35 autoimmune/inflammatory diseases and 27 immune-related quantitative traits), we found that 89 of the 123 studies incorporate three or more different genotyping arrays (**Supplementary Tables 1 and 2**).

Because genotyping arrays and imputation panels can vary substantially, SNP coverage is highly heterogeneous, ranging from hundreds of thousands to tens of millions of variants (**Fig. 2a**). Across the same group of 123 autoimmune GWAS, the smallest GWAS examined 520,268 variants, whereas the largest included 56,839,225 variants — a more than 100-fold difference in variants profiled (**Fig. 2b and Methods**). This variability presents significant challenges for downstream analyses of GWAS summary statistics, particularly methods that require intersections of variants across multiple studies, such as multi-trait colocalization^24^ and GWAS meta-analyses^25^, where intersecting such disparate SNP sets results in substantial data loss. Additionally, GWAS summary statistics may be released in different genome builds, requiring liftover to a common reference (**Fig. 2b**). In practice, GWAS metadata is not always clearly reported, and substantial manual effort is often required to infer the genome build and harmonize variant identifiers, reference and alternate alleles, and effect allele annotations.

**Fig. 2.**
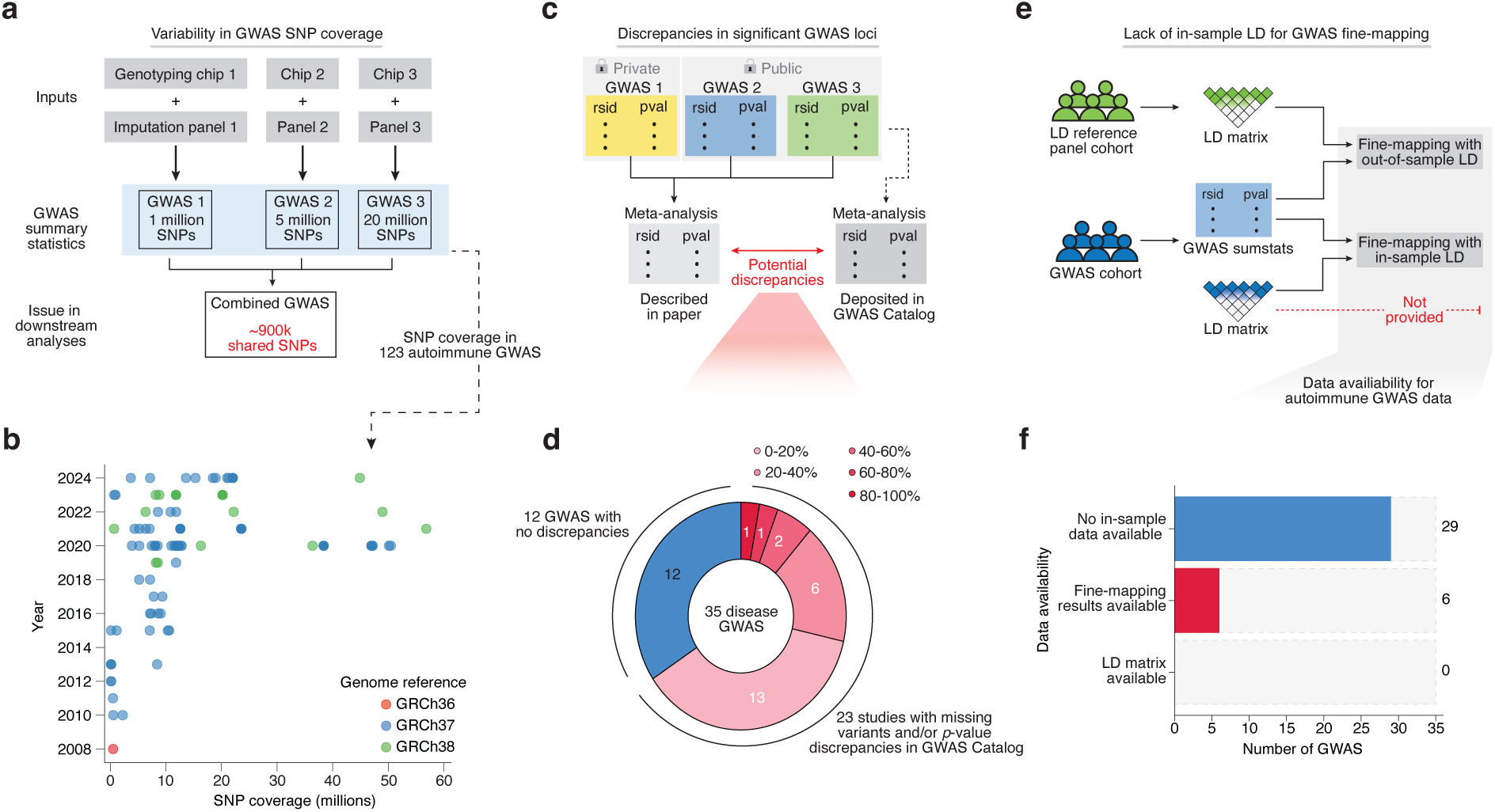
Limitations of GWAS and data sharing can hinder identification of putative causal variants. **a,** GWAS SNP coverage is determined by the genotyping array and imputation panel used by the authors. A common step in downstream analyses utilizing multiple GWAS is taking the intersection of the SNP sets which can lead to large data losses due to the variability in genotyping technologies and reference panels used. **b,** SNP coverage for 123 autoimmune GWAS studies ranging from 2008 to 2024 showcase the wide variability in SNP coverage along with the genome-build that the published data is released on. **c,** Discrepancies can occur between the significant GWAS loci reported by the authors in their paper and the data released to the public due to the inclusion of private datasets in GWAS meta-analyses. These discrepancies typically take the form of differences in p-values (where a variant goes from being genome-wide significant to no longer being so), or less often, a reported lead variant for a locus does not occur in the publicly released summary statistics. **d,** Across the largest GWAS for each unique disease in our GWAS dataset, 23 out of 35 GWAS contained at least 1 discrepancy. These 23 studies are further categorized by the percentage of author reported loci that contained a discrepancy. **e,** A common limitation with GWAS is that the authors typically do not release in-sample LD matrices or fine-mapping results using the in-sample LD. This forces downstream researchers to utilize out-of-sample reference panels for fine-mapping. **f,** Again utilizing the largest GWAS for each unique disease, we observed 29 out of 35 GWASs did not provide any in-sample LD data, while 6 studies provided fine-mapping results using in-sample LD.

Discrepancies can also arise between variants reported as significant by study authors and those present in publicly available GWAS summary statistics. Due to a combination of private and public meta-analyses, authors may have access to datasets that are not included in released summary statistics (**Fig. 2c**). Consequently, reported lead variants may not reach genome-wide significance in the released summary statistics. When examining the largest GWAS available for each of 35 autoimmune diseases (**Supplementary Table 3**), we found that 23 studies exhibited discrepancies where a reported significant GWAS variant is not present or has a non-significant *p*-value in the publicly released summary statistics (**Fig. 2d**).

A common task after GWAS — necessary for any experiments targeting GWAS loci — is fine-mapping, i.e. statistical modeling to distinguish the likely causal genetic variants driving the association signal in a given locus from non-causal variants that are associated merely because of LD with the true causal variant. High-quality fine-mapping relies on GWAS summary statistics and an LD matrix from the same cohort used in the GWAS or from a closely-matched population, because population differences in LD structure reduce the quality of fine-mapping analyses^26,27^ (**Fig. 2e**). However, in-sample LD matrices are rarely provided with the GWAS summary statistics: Across the 35 autoimmune disease GWAS, none of the studies provided an in-sample LD matrix and only six studies reported fine-mapping using in-sample LD (**Fig. 2f**). For the 29 studies without any fine-mapping results or to use newer fine-mapping methods, it is necessary to use out-of-sample LD matrices with the potential for LD mismatch, as we detail in the next section. While sharing of LD matrices has challenges related to data privacy and very large file sizes, improved practices for LD data sharing would greatly enhance fine-mapping efforts.

### LD dropout is prevalent across GWAS and hinders statistical fine-mapping

In statistical fine-mapping, the first goal is to group variants into credible sets with a defined probability of containing the true causal variant. Accurate identification of credible sets depends heavily on minimizing mismatch between the variants genotyped in the GWAS and those included in the LD reference panel.

In addition to LD mismatch between populations, another seemingly trivial but highly consequential and poorly characterized issue with out-of-sample LD reference panels is LD dropout, which is defined as the proportion of GWAS variants that are absent from the LD reference panel and thus excluded from fine-mapping. Using the PanUKBB LD dataset from 441,331 total individuals, we found that the LD dropout varies widely across the 123 autoimmune GWAS, ranging from full coverage (0% dropout) to nearly 60% dropout (**Fig. 3a**). Intuitively, LD dropout should be more severe for less powered ancestries due to lower quality reference panels but this is not always the case. We found that studies in non-European ancestries experienced higher dropout rates, while, surprisingly, studies in European ancestries can also experience substantial LD dropout depending on the study. The latter was most often due to the genotyping technology used in the GWAS, such as customized genotyping chips or whole genome sequencing. To understand whether the LD dropout varies across GWAS loci within a particular study, we computed the average and maximum amount of dropout across all autoimmune GWAS loci in 1 Mbp windows using the PanUKBB LD panel (**Fig. 3b**). We find that LD dropout can non-uniformly affect loci: For example, in a recently published inflammatory bowel disease GWAS^28^, we find certain loci that have high dropout only within specific sub-regions and other loci with more uniform dropout across the fine-mapping window.

**Fig. 3.**
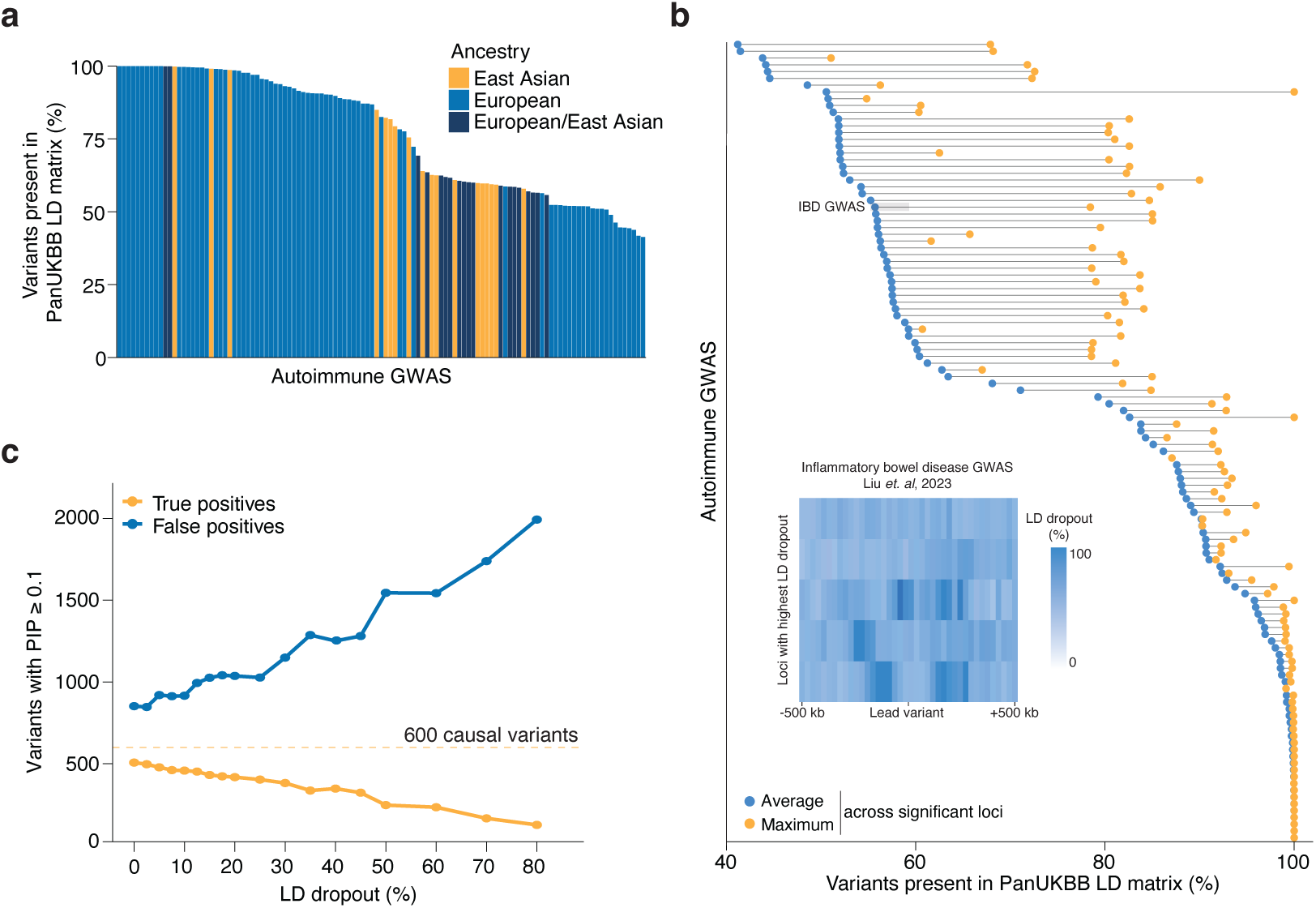
Linkage disequilibrium (LD) matrix variant dropout is widespread and leads to nomination of false positives after fine-mapping. **a,** Percentage of variants present in the ancestry matched PanUKBB LD matrix for each of the 123 autoimmune GWAS. **b,** Average and maximum percentage of variants present at a locus level across each of the GWAS. *Inset:* Spatial distribution of LD dropout at 5 inflammatory bowel disease GWAS loci. **c,** Total count of true positive (TP) and false positive (FP) variants across 300 simulated GWAS loci across a wide range of LD dropout. Each simulated GWAS locus contained 2 causal variants (*n* = 600 causal variants total), where TPs are defined as causal variants given a posterior inclusion probability (PIP) greater than or equal to 0.1, and FPs are defined as non-causal variants given a PIP greater than or equal to 0.1.

To better characterize how LD dropout can impact causal variant identification and fine-mapping accuracy, we simulated GWAS summary statistics for 300 GWAS loci with 2 causal variants each, and modeled different levels of variant dropout^29^ (**Methods**). We then performed fine-mapping using Sum of Single Effects regression (SuSiE)^30^ with the simulated summary statistics and ancestry-matched LD matrix from PanUKBB^31^. Non-causal variants with a posterior inclusion probability (PIP) greater than the commonly used threshold of 0.1 were assigned as false positives^32,33^. As LD dropout increases, false positives increase and true positives decrease (**Fig. 3c**). Even without any dropout, there are a substantial number of false positives — a notable limitation of statistical fine-mapping. Altogether, these results from existing autoimmune GWAS and simulated GWAS with ground-truth causal variants highlight the challenges for nominating likely causal variants given the lack of shared LD data. New ways to share LD data that address privacy concerns and that efficiently reduce the typically large file sizes of LD matrices will be necessary to improve fine-mapping efforts.

### Incorporating data across traits and modalities can increase power for variant nomination

The increasingly deep GWAS data from multiple related traits provide an opportunity to analyze their shared genetic architecture and mechanism across traits/diseases^34–36^. At the locus level, colocalization of multiple putatively shared signals can identify shared causal variants, leveraging power across GWAS studies without the need for LD matrices^24^. This approach can also nominate a given locus and its fine-mapped variants for a disease/trait where the association signal does not reach genome-wide significance. Across 123 autoimmune GWAS, we performed multi-trait colocalization with HyPrColoc to nominate variants in loci containing signals shared across multiple traits^24^. As an example, we colocalized a GWAS signal across 16 autoimmune-related GWAS (9 unique diseases) in a 200 kb region centered on chromosome 4 (**Supplementary Table 4**). Six out of the 16 GWAS contained at least one genome-wide significant variant in that window. HyPrColoc identified a single variant (chr4:40,307,564) with a large posterior probability that appears to explain the signal underlying 16 colocalized GWAS (**Fig. 4a**). This illustrates how shared genetic architecture of related traits allows identification of more high-confidence variants.

**Fig. 4.**
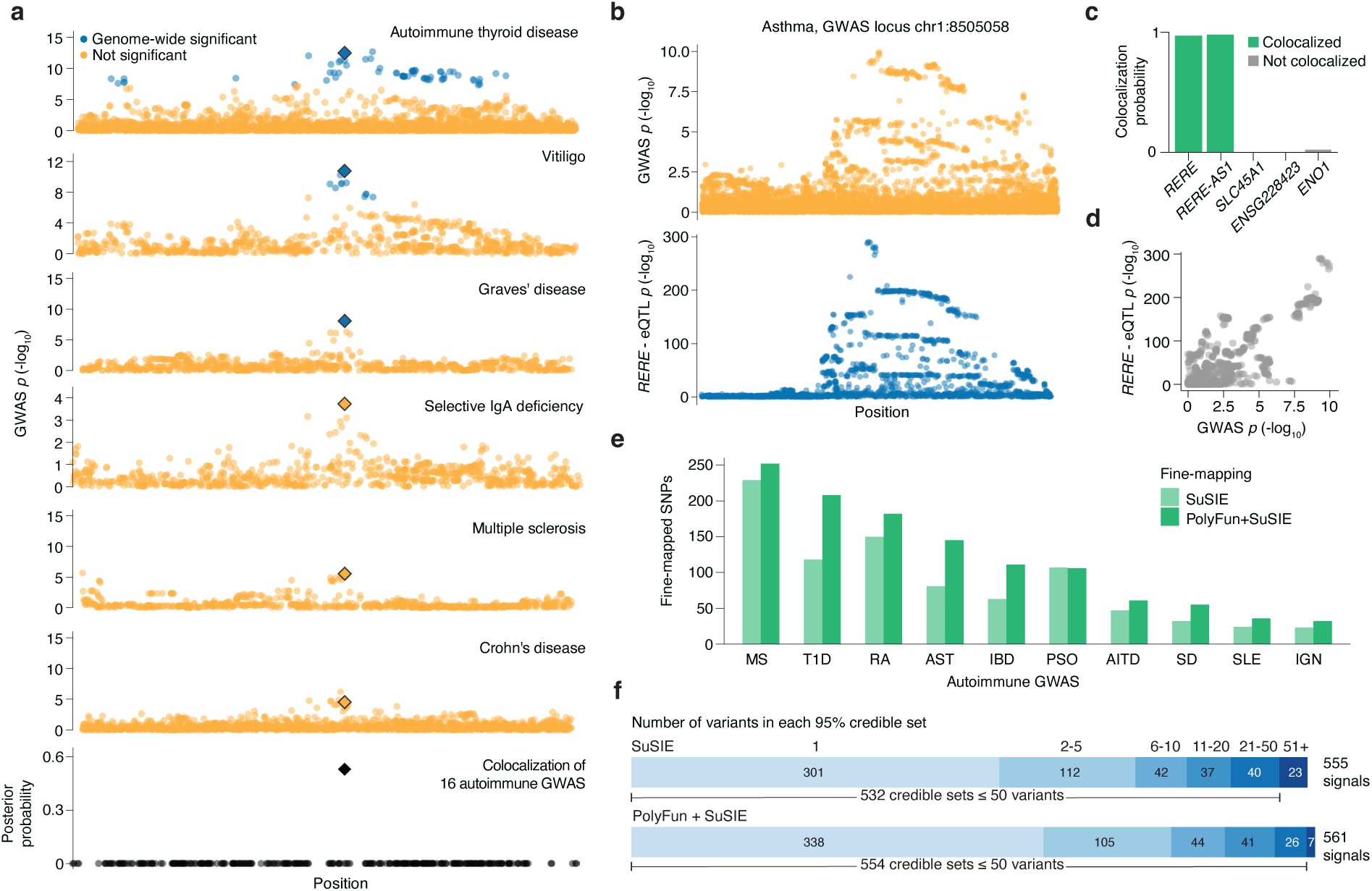
Colocalization with multiple GWAS, molecular quantitative trait loci (molQTLs) and functional genomics improves identification of putative causal variants. **a,** Multi-trait colocalization across 16 autoimmune GWAS (9 unique diseases) using HyPrColoc identifies a single variant (rs13136820) with a relatively high posterior probability. This colocalization included 10 GWAS where there were no genome-wide significant variants in this locus, showcasing the ability of colocalization to discover signal in less-powered GWAS (**Supplementary Table 4**). **b-d,** An example locus from an Asthma GWAS where we attempt to colocalize it with nearby genes using the INTERVAL eQTL dataset^39^. We identified two significant colocalizations with the gene RERE and its antisense copy providing a potential molecular explanation for the GWAS locus. **e,** Functional fine-mapping with PolyFun across 250 GWAS loci in 10 unique autoimmune diseases (MS: Multiple sclerosis, T1D: Type 1 diabetes, RA: Rheumatoid arthritis, AST: Asthma, IBD: Inflammatory bowel disease, PSO: Psoriasis, AITD: Autoimmune thyroid disease, SD: Sjögren’s disease, SLE: Systemic lupus erythematosus, IGN: IgA nephropathy) identifies 36% more variants with a PIP above 0.1. **f**, Number of variants in 95% credible sets when fine-mapping the same 250 GWAS loci from **(e)** using SuSIE with and without PolyFun. 338 signals were fine-mapped to a single variant when utilizing PolyFun compared to only 301 signals using only SuSIE.

GWAS loci can also be colocalized with genetic associations to molecular traits, such as molecular quantitative trait loci (molQTLs), e.g., for expression (eQTL) and chromatin accessibility (caQTL)^37,38^. Similar to multi-trait GWAS colocalization, these methods can refine association signals to improve fine-mapping, and further pinpoint putative molecular changes, such as gene expression changes. To illustrate this approach, we performed GWAS–eQTL colocalization at an asthma-associated locus using the INTERVAL eQTL dataset^39^. At one locus, we observe strong evidence of colocalization for the gene *RERE* and the long noncoding RNA *RERE-AS1* using coloc^40^ (**Fig. 4b-d**). Notably, a recent study implicated *RERE* as contributing to bronchial asthma^41^. Chromatin accessibility QTLs have even higher GWAS colocalization rates, linking GWAS variants with changes in the chromatin state^42^. While GWAS-molQTL colocalization has its shortcomings such as biases in identification of true causal genes^43^ often lacks gene expression data from the most relevant cell type^44^, it provides an additional, independent layer of information for identifying putative causal variants and their molecular mechanisms.

Integrating GWAS data with biochemical hallmarks of regulatory elements can better localize causal genetic variants and pinpoint relevant cell types where variants may play a role in gene regulation^45,46^. In addition to assessing overlap between putative causal variants and these biochemical hallmarks in specific loci, multiple statistical approaches have been developed to directly incorporate these data in fine-mapping. One recent method from Price and colleagues (PolyFun)^16^ annotates variants based on their functional role (e.g. untranslated regions, enhancers, introns, promoters, etc.), conservation score and population genetic data to inform priors on variant causality^47^, using in total more than 150 features. To assess the impact of incorporating such annotations on variant prioritization, we fine-mapped 250 loci across 10 autoimmune diseases from 26 distinct GWAS using PolyFun. We found that across these loci, this approach identified more variants with PIPs above 0.1 in comparison to statistical fine-mapping alone (using SuSiE) (**Fig. 4e**). Across 250 loci, functionally-informed fine-mapping identified 1,188 SNPs with a PIP above 0.1 (36% increase), while SuSiE alone identified 874 SNPs with a PIP above 0.1. When used to inform priors, functional annotations can improve fine-mapping resolution: Across 250 GWAS loci, PolyFun resolved 338 signals to a single variant, while SuSiE only resolved 301 signals to a single variant (**Fig. 4f**). Thus, using functional annotation to inform fine-mapping can capture more variants and improve the signal-to-noise ratio in GWAS loci.

### Emerging deep learning models for variant prioritization

Convolutional neural networks with transformer architectures have had a major impact as general artificial intelligence systems. Recently, similar models have been built to predict the functional impact of non-coding variants, although rigorous testing with large datasets of known functional variants has been limited. These sequence-to-function methods have the potential to transform GWAS interpretation, as they allow querying any possible variant’s allelic effects on chromatin or gene expression, sometimes in a cell-type specific manner^48–50^. Here, we investigated the performance of multiple recent models for prioritizing likely functional variants from GWAS loci validated through CRISPR screening — one of the first systematic evaluations of its kind.

We focus our analysis on blood trait GWAS loci that we recently experimentally characterized in K562 cells by STING-seq, an approach that combines statistical fine-mapping, CRISPR interference (CRISPRi) targeting of variant positions in candidate *cis*-regulatory elements (cCRE) and single cell RNA-seq (scRNA-seq)^51^ (**Fig. 5a**). Out of the targeted 543 variant positions, 136 had a significant association to at least one nearby gene (here called STING-positives), and 407 variants with no association (STING-negatives). We note that some of the STING-negatives are likely false negatives due to lack of power in the sparse scRNA-seq data^52^. Furthermore, for STING-positives, we do not know explicitly that the variant itself is causal since precise editing was not performed for all variants.

**Fig. 5.**
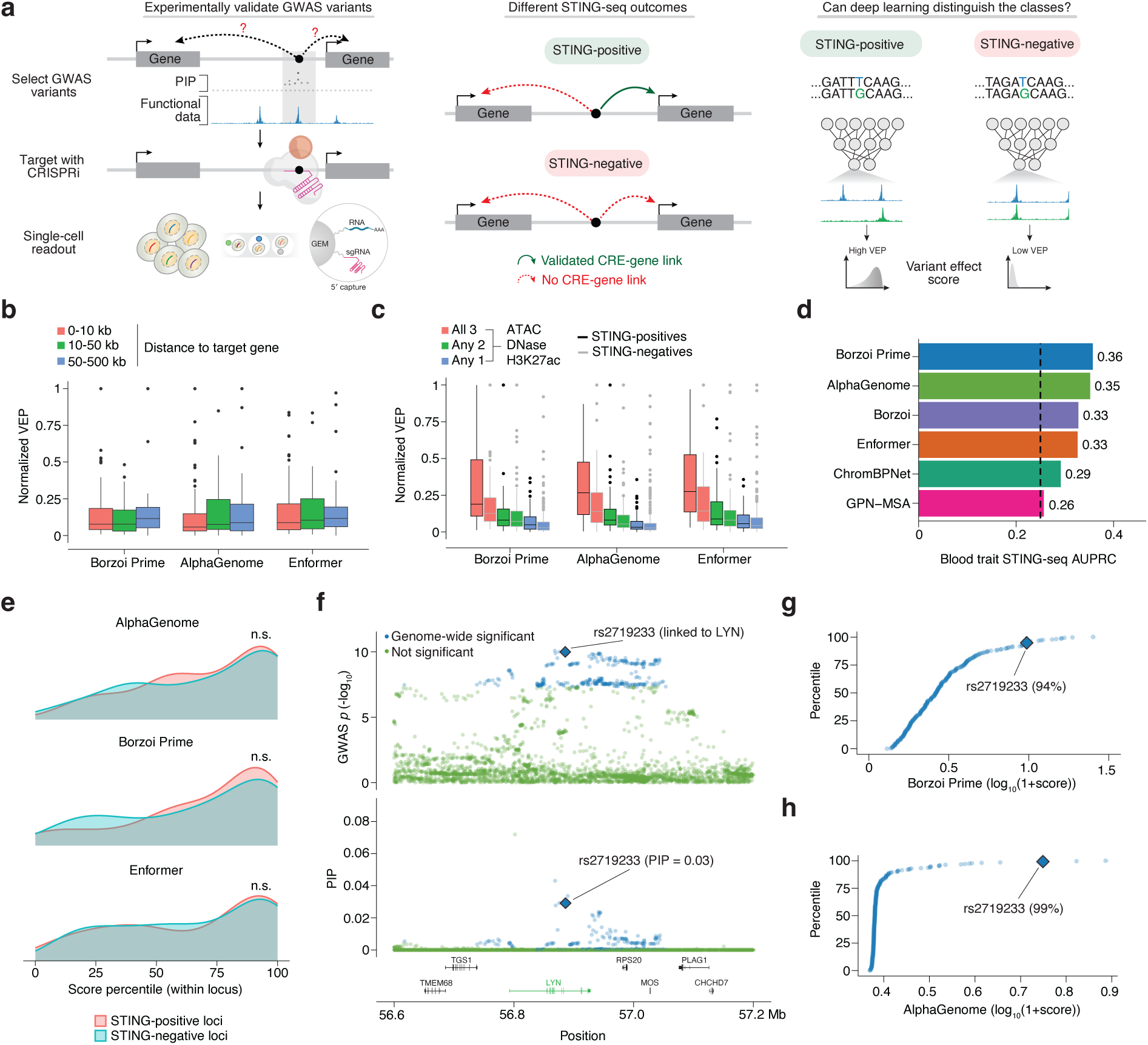
Deep learning variant effect predictors show promise in variant prioritization with room for improvement. **a,** Morris et al. (2023) performed a scCRISPRi screen targeting GWAS variants from blood traits. They identified 136 non-coding variants regulating nearby genes (STING-positives), along with 407 non-coding variants that do not appear to regulate nearby genes (STING-negatives). We utilize this experimentally validated dataset to assess the ability of deep learning variant effect predictors in prioritizing truly causal variants. **b,** Stratifying the variant effect scores for the STING-positives based on their distances to their true target gene. **c,** Stratifying the variant effect scores for both STING-positives and -negatives based on the number of functional annotations (ATAC, DNase, H3K27ac) each variant possessed. **d,** Borzoi Prime and AlphaGenome yielded the largest AUPRCs when we applied each tool to the binary classification task of distinguishing between STING-positives and -negatives. The baseline performance for a random classifier in this task is 0.25, visualized by the vertical dashed line. For scoring, we utilized the same approach used by a recent benchmarking study^75^ (**Methods**). **e,** Percentile for each STING-positive and -negative variants based on the variant effect score when compared to all the genome-wide significant variants in their respective locus. This highlights that both classes of variants were enriched in the top quartile of variant effect scores within their locus. Differences in variant effect scores between STING-positive and -negative variants were not significant (Mann-Whitney *U* test, *p* > 0.05). **f,** GWAS locus for monocyte percentage which contains a STING-positive variant, rs2719233, identified by Morris et al. (2023) to regulate the gene *LYN*. This variant was given a relatively low PIP (0.029) indicating that fine-mapping struggled in this locus to identify variants with high confidence. **g-h,** Variant effect scores for all genome-wide significant variants in the example locus from **(f)**. The STING-positive variant (rs2719233) occurs in the 94^th^ and 99^th^ percentile for Borzoi Prime and AlphaGenome respectively showcasing the ability to use deep learning predictors in loci where fine-mapping struggles.

We obtained variant prediction scores from sequence-to-function and conservation-based models: Enformer^53^, Borzoi^49^, Borzoi Prime^50^, GPN-MSA^54^, ChromBPNet^55^ and AlphaGenome^48^. First, we investigated how the predicted variant effects correlate with different features. Variant scores had little correlation to distance to the STING-seq target gene of STING-positives (**Fig. 5b**), but variants with more functional marks had higher scores across both variant sets (**Fig. 5c**). Next, we compared the model’s ability to distinguish STING-positive and STING-negative variants. All of the models outperform baseline (random) — even ChromBPNet, that, as a chromatin model, is not designed for transcription prediction. Borzoi Prime and AlphaGenome have the highest AUPRCs but only moderately above the baseline level (**Fig. 5d and Supplementary Fig. 1**). This may be because both STING-positive and -negative variants have functional annotations, making the classification task particularly difficult.

To analyze the performance of deep learning models for variant prioritization within a locus, we computed AlphaGenome, Borzoi Prime, and Enformer scores for all genome-wide significant GWAS variants. Next, we selected 50 loci containing STING-positive variants and 50 loci containing STING-negative variants (**Supplementary Fig. 2**). We found that the variants targeted in the STING-seq study were enriched in the top quartiles of all GWAS variants, suggesting that taking the top *n* variants could be a reasonable strategy for prioritizing GWAS variants. However, the enrichment was similar for both STING-positive and -negative variants (*p* > 0.05), indicating that the models may simply prioritize variants in functional chromatin contexts, rather than being able to predict their eventual signal in the genetic perturbation assay (**Fig. 5e**).

To better understand whether deep learning models can identify causal variants, we examined a locus for a quantiative blood trait (monocyte percentage) where silencing the variant rs2719233, which is in a later intron of *LYN*, affected the expression of *LYN* (**Fig. 5f**). This variant is in tight LD with a number of other variants, leading to a large credible set even after fine-mapping. Morris et al. (2023) targeted rs2719233 due to its overlap with H3K27ac ChIP-seq data from K562 cells. We scored all 300 significant GWAS variants in this locus with Borzoi Prime (**Fig. 5g**) and AlphaGenome (**Fig. 5h**) and observed that rs2719233 was in the 94th and 99th percentile, respectively, indicating good prioritization of this variant above other variants with significant associations. This is an example for how deep learning models can complement more traditional statistical genetics methods for prioritizing variants for functional follow-up; however, as larger perturbation-based datasets become available from individual labs, industry and larger consortia, it will be vital to carefully evaluate model performance through established predictive measures such as precision and recall.

### How do we make the most of GWAS data resources?

The quality and completeness of GWAS summary statistics are crucial determinants of many downstream analyses, including the ultimate translation of genetic associations into biological mechanisms, biomarkers and therapeutic targets. We highlight pervasive bottlenecks in the post-GWAS discovery pipeline: inconsistencies in SNP coverage and imputation, and the frequent absence of in-sample resources such as LD matrices and fine-mapping results. Reliance on out-of-sample LD reference panels introduces the subtle yet consequential problem of LD dropout, in which differences between the GWAS and reference panel cohorts distort downstream analysis. Such mismatches can artificially inflate PIPs for non-causal variants, which can squander precious resources for functional follow-up efforts on false positives. Improving variant prioritization and the efficiency of functional validation will require in-sample LD resources and well-annotated, harmonized summary statistics.

While logistical barriers have historically limited the release of in-sample LD resources, these challenges are not insurmountable. A fundamental shift in how LD data are generated, stored, and disseminated is warranted. The size of LD matrices poses constraints in cost-effective data sharing, however, methodological advancements have enabled efficient compression^56^ or construction of sparse LD matrices without substantial loss of accuracy^57^. Concerns regarding preservation of individual-level genotype privacy, especially in meta-analytic contexts, have similarly been addressed through the generation of pseudo-genotypes that replicate in-sample LD structure while discarding individual-level genetic content^58^. Cloud-based analysis platforms such as the UK Biobank Research Analysis Platform (UKB-RAP)^59^ and the All of Us Researcher Workbench^60^ offer an alternative model, providing researchers remote access to in-sample data. With increasing access to in-sample LD information, continued harmonization efforts, such as the GWAS Summary Statistics Format (GWAS-SSF) from the NHGRI-EBI GWAS Catalog^1^, will be imperative. The inclusion of the standardized data fields necessary for full downstream utilization would increase the value of future GWAS efforts.

The evolution of fine-mapping is being driven by increasingly robust genomic datasets and integrative methodological developments. Greater diversity in GWAS cohorts will facilitate causal variant localization by leveraging differences in LD structures across ancestries^61^. Further, the development of high-quality ancestry-specific reference panels, such as PanUKBB^31^ and All of Us^60^, is expected to refine large European LD blocks through multi-ancestry fine-mapping. The ongoing shift from SNP array based GWAS toward whole-genome sequencing eliminates the reliance on imputation and enables more accurate characterization of LD structure. The recent release of whole-genome sequencing data for 490,640 UK Biobank participants is a prime example of this transition^62^. Concurrently, methodological developments in Bayesian fine-mapping aim to improve resolution through incorporation of informative priors, including multi-ancestry GWAS data^63–65^, functional genomic annotations^45,66,67^, and deep learning variant effect predictions^68^.

Improving the predictive modeling of variant effects will require systematic generation of perturbation datasets in training and testing data, rather than simply relying on observed variants. Initiatives such as Impact of Genomic Variation on Function (IGVF) consortium aim to comprehensively perturb candidate variants and document their regulatory effect^69^. Recent studies have applied high-throughput screening technologies such as CRISPRi or MPRA to study regulatory variants in GWAS loci^70–72^. Increasing context-specific perturbation datasets will further assist in fine-tuning deep learning models through causal refinement, in which targeted perturbations that expose missing regulatory mechanisms are incorporated to improve predictive performance^73^. Critically, null perturbations within these catalogues provide essential negative examples for model training, facilitating differentiation between causal variants and surrounding linkage noise.

The field must move beyond the cataloguing of genetic associations toward the thorough interrogation of underlying causal mechanisms. Realizing the full potential of GWAS will require concerted efforts on multiple fronts beyond simply increasing the size of GWAS data sets: improved data sharing, continued methodological development, and expansion of perturbation datasets. These developments will trigger a positive feedback cycle in which higher-quality data will enable more accurate variant effect predictions, thereby guiding experimental efforts toward efficient experimental interrogation and perturbation in relevant cellular contexts. Ultimately, translating genetic associations into clinical insights will require the field to unite around rigorous, inclusive, and multi-dimensional frameworks for post-GWAS discovery.

## Methods

### GWAS dataset curation

For the autoimmune GWAS dataset, we extracted summary statistics from 123 GWAS covering a spectrum of autoimmune diseases. We only included GWAS whose publicly available summary statistics included significance, effect sizes, and standard errors, which are required to perform fine-mapping.

### Fine-mapping with SuSiE

SuSiE was run using the R package susieR with the the susie_rss function^14^. Betas and standard errors were obtained from the GWAS summary statistics, and the LD matrix used for fine-mapping was ancestry-matched with the GWAS from PanUKBB^31^. When running SuSiE, we set *L* = 10 which allows for up to 10 independent signals to be identified in the locus.

### GWAS and LD dropout simulation

GWAS summary statistics were simulated using the tool GWASBrewer^74^. The pi parameter was set to 2/N (N being the total number of variants in the locus), and pi_exact was set to TRUE in order to ensure there would be exactly two causal variants per locus. For different GWAS parameters such as sample size (500,000) and heritability (0.03), we utilized realistic values informed by previous simulation studies^27^. We used PanUKBB LD matrices from 10 different autoimmune GWAS loci^31^. In total, we simulated 300 GWAS loci that were used for our analysis of LD dropout.

For each dropout percentage, we randomly selected a group of variants in the GWAS summary statistics to remove to simulate LD dropout. We removed the corresponding rows from the LD matrix. Next, we fine-mapped the GWAS using SuSIE and then quantified the number of true positives and false positives. True positives are defined as truly causal variants that received a PIP of 0.1 or greater. False positives are defined as variants that are not truly causal and obtain a PIP of 0.1 or greater.

### Functional fine-mapping with PolyFun

We selected 250 random autoimmune GWAS loci and fine-mapped using PolyFun^67^. We utilized the precomputed prior probabilities provided by the authors before running the PolyFun fine-mapping script which is a wrapper around SuSiE.

### Multi-trait colocalization with HyPrColoc

We utilized HyPrColoc to identify regions in the genome with shared signals across multiple studies^24^. For each region, we intersected the GWAS summary statistics across all studies to identify shared variants, requiring a minimum of 200 variants in the region prior to running HyPrColoc.

HyPrColoc outputs subsets of the input GWAS set that colocalize together with a certain cluster posterior probability. For each subset, HyPrColoc outputs a PIP for each variant. We computed the posterior probability of each SNP being causal in a specific region by multiplying the PIP with the cluster posterior probability.

### GWAS-eQTL colocalization with Coloc

We utilized the INTERVAL dataset^39^ of eQTLs from blood cells and ran the analyses using coloc^40^. For each locus, we identified all protein-coding and lncRNA genes within 1Mb from the lead SNP and attempted to colocalize the GWAS locus with eQTLs of each gene. We used a threshold of 0.5 on the H4 probability to determine if a GWAS locus colocalized with an eQTL.

### Deep learning variant effect prediction

We used multiple deep learning models for scoring variant effects: Enformer^53^, Borzoi^49^, Borzoi Prime^50^, GPN-MSA^54^, ChromBPNet^55^ and AlphaGenome^48^. For scoring, we focused our analyses on SNPs since indels could lead to a shifted context which makes proper scoring of variants non-trivial. We followed the approach used by Benegas et al. in their TraitGym paper for scoring SNPs^75^. For Enformer, Borzoi, and Borzoi Prime, we used the “L2 of L2 norms” score described by Benegas et al. For AlphaGenome, we took the maximum value across the set of L2 norms as recommended by the AlphaGenome authors^48^. For ChromBPNet, we utilize the K562 model published with the paper and the variant-scorer codebase (https://github.com/kundajelab/variant-scorer/tree/main) where we use the abs_logfc score. Lastly for GPN-MSA^54^, we utilize the pre-computed scores for all possible SNPs in hg38 (https://huggingface.co/datasets/songlab/gpn-msa-hg38-scores).

## Data availability

Data sources for all GWAS utilized in this study are listed in *Supplementary Table 1*. The GWAS and target genes for deep learning variant effect prediction were obtained from Morris et al. (2023).

## Code availability

Source code is provided via GitHub at https://github.com/oma219/GWASperspective.

## Acknowledgements

We thank the Sanjana, Lappalainen, and Singh laboratories for support and advice. D.S. and S.G. are supported by the Janeway Fellowship. T.S. is supported by John and Wendy Havens, the Columbia Precision Medicine Initiative, and a NARSAD Young Investigator Grant. T.L. is supported by the NIH (R01MH106842, 24HG012090, R01HL168247 and R01HG012790), the European Research Council (ERC) under the European Union’s Horizon 2020 Research and Innovation Program (101043238), and Knut and Alice Wallenberg Foundation grant KAW 2023.0337. N.E.S. is supported by the NIH/NHGRI (DP2HG010099 and R01HG012790), the NIH/National Cancer Institute (NCI) (R01CA279135 and R01CA218668), the NIH/National Institute of Allergy and Infectious Diseases (NIAID) (R01AI176601), the NIH/National Heart, Lung, and Blood Institute (NHLBI) (R01HL168247), the Simons Foundation for Autism Research, the MacMillan Center for the Study of the Noncoding Cancer Genome, and New York University and New York Genome Center funds.

## Contributions

Conceived and designed the study: O.Y.A., N.S., T.L., N.E.S. Collected and integrated GWAS datasets: O.Y.A., N.S., D.S. Performed analysis: O.Y.A., N.S., A.B.R., D.S., A.D., S.G. Supervised the research: T.S., T.L., N.E.S. Wrote the paper: O.Y.A., N.S., T.L., N.E.S. Edited and approved the final manuscript: all authors

## Competing interests

N.E.S. is an adviser to Qiagen and is a cofounder and adviser of OverT Bio and TruEdit Bio. T.L. is an adviser to Goldfinch Bio and GSK and, with equity, Variant Bio.

## References

1. Cerezo, M. et al. The NHGRI-EBI GWAS Catalog: standards for reusability, sustainability and diversity. Nucleic Acids Res 53, D998–D1005 (2025).

2. Claussnitzer, M. et al. FTO Obesity Variant Circuitry and Adipocyte Browning in Humans. New England Journal of Medicine 373, 895–907 (2015).

3. Minikel, E. V., Painter, J. L., Dong, C. C. & Nelson, M. R. Refining the impact of genetic evidence on clinical success. Nature 629, 624–629 (2024).

4. Thein, S. L., Menzel, S., Lathrop, M. & Garner, C. Control of fetal hemoglobin: new insights emerging from genomics and clinical implications. Hum Mol Genet 18, R216–223 (2009).

5. Bauer, D. E. et al. An erythroid enhancer of BCL11A subject to genetic variation determines fetal hemoglobin level. Science 342, 253–257 (2013).

6. Canver, M. C. et al. BCL11A enhancer dissection by Cas9-mediated in situ saturating mutagenesis. Nature 527, 192–197 (2015).

7. Frangoul, H. et al. CRISPR-Cas9 Gene Editing for Sickle Cell Disease and β-Thalassemia. New England Journal of Medicine 384, 252–260 (2021).

8. Maurano, M. T. et al. Systematic Localization of Common Disease-Associated Variation in Regulatory DNA. Science 337, 1190–1195 (2012).

9. Adzhubei, I., Jordan, D. M. & Sunyaev, S. R. Predicting Functional Effect of Human Missense Mutations Using PolyPhen-2. Curr Protoc Hum Genet 0 7, Unit7.20 (2013).

10. Cheng, J. et al. Accurate proteome-wide missense variant effect prediction with AlphaMissense. Science 381, eadg7492 (2023).

11. Rentzsch, P., Witten, D., Cooper, G. M., Shendure, J. & Kircher, M. CADD: predicting the deleteriousness of variants throughout the human genome. Nucleic Acids Res 47, D886–D894 (2019).

12. Abell, N. S. et al. Multiple causal variants underlie genetic associations in humans. Science 375, 1247–1254 (2022).

13. Long, E. et al. Massively parallel reporter assays and variant scoring identified functional variants and target genes for melanoma loci and highlighted cell-type specificity. Am J Hum Genet 109, 2210–2229 (2022).

14. Wang, G., Sarkar, A., Carbonetto, P. & Stephens, M. A Simple New Approach to Variable Selection in Regression, with Application to Genetic Fine Mapping. J. R. Stat. Soc. Ser. B. Stat. Methodol. 82, 1273–1300 (2020).

15. Amariuta, T. et al. Improving the trans-ancestry portability of polygenic risk scores by prioritizing variants in predicted cell-type-specific regulatory elements. Nat Genet 52, 1346–1354 (2020).

16. Weissbrod, O. et al. Leveraging fine-mapping and multipopulation training data to improve cross-population polygenic risk scores. Nat Genet 54, 450–458 (2022).

17. Huerta-Sánchez, E. et al. Altitude adaptation in Tibetans caused by introgression of Denisovan-like DNA. Nature 512, 194–197 (2014).

18. Sabeti, P. C. et al. Genome-wide detection and characterization of positive selection in human populations. Nature 449, 913–918 (2007).

19. Marbach, D. et al. Tissue-specific regulatory circuits reveal variable modular perturbations across complex diseases. Nat Methods 13, 366–370 (2016).

20. Reshef, Y. A. et al. Detecting genome-wide directional effects of transcription factor binding on polygenic disease risk. Nat Genet 50, 1483–1493 (2018).

21. Marchini, J. & Howie, B. Genotype imputation for genome-wide association studies. Nat Rev Genet 11, 499–511 (2010).

22. Visscher, P. M. et al. 10 Years of GWAS Discovery: Biology, Function, and Translation. Am J Hum Genet 101, 5–22 (2017).

23. Cortes, A. & Brown, M. A. Promise and pitfalls of the Immunochip. Arthritis Res Ther 13, 101 (2011).

24. Foley, C. N. et al. A fast and efficient colocalization algorithm for identifying shared genetic risk factors across multiple traits. Nat Commun 12, 764 (2021).

25. Willer, C. J., Li, Y. & Abecasis, G. R. METAL: fast and efficient meta-analysis of genomewide association scans. Bioinformatics 26, 2190–2191 (2010).

26. Chen, W. et al. Improved analyses of GWAS summary statistics by reducing data heterogeneity and errors. Nat Commun 12, 7117 (2021).

27. Kanai, M. et al. Meta-analysis fine-mapping is often miscalibrated at single-variant resolution. Cell Genom 2, 100210 (2022).

28. Liu, Z. et al. Genetic architecture of the inflammatory bowel diseases across East Asian and European ancestries. Nat Genet 55, 796–806 (2023).

29. Morrison, J. GWASBrewer: An R Package for Simulating Realistic GWAS Summary Statistics. Genet Epidemiol 49, e22594 (2025).

30. Zou, Y., Carbonetto, P., Wang, G. & Stephens, M. Fine-mapping from summary data with the “Sum of Single Effects” model. PLOS Genetics 18, e1010299 (2022).

31. Karczewski, K. J. et al. Pan-UK Biobank genome-wide association analyses enhance discovery and resolution of ancestry-enriched effects. Nat Genet 57, 2408–2417 (2025).

32. Cai, M. et al. XMAP: Cross-population fine-mapping by leveraging genetic diversity and accounting for confounding bias. Nat Commun 14, 6870 (2023).

33. Wang, J. et al. Fine-mapping methods for complex traits: essential adaptations for samples of related individuals. Brief Bioinform 26, bbaf614 (2025).

34. Bulik-Sullivan, B. K. et al. LD Score regression distinguishes confounding from polygenicity in genome-wide association studies. Nat Genet 47, 291–295 (2015).

35. Demela, P., Pirastu, N. & Soskic, B. Cross-disorder genetic analysis of immune diseases reveals distinct gene associations that converge on common pathways. Nat Commun 14, 2743 (2023).

36. Lincoln, M. R. et al. Genetic mapping across autoimmune diseases reveals shared associations and mechanisms. Nat Genet 56, 838–845 (2024).

37. Aguet, F. et al. Molecular quantitative trait loci. Nat Rev Methods Primers 3, 4 (2023).

38. GTEx Consortium. The GTEx Consortium atlas of genetic regulatory effects across human tissues. Science 369, 1318–1330 (2020).

39. Tokolyi, A. et al. The contribution of genetic determinants of blood gene expression and splicing to molecular phenotypes and health outcomes. Nat Genet 57, 616–625 (2025).

40. Giambartolomei, C. et al. Bayesian test for colocalisation between pairs of genetic association studies using summary statistics. PLoS Genet 10, e1004383 (2014).

41. Zaied, R. E., Gokuladhas, S., Walker, C. & O’Sullivan, J. M. Unspecified asthma, childhood-onset, and adult-onset asthma have different causal genes: a Mendelian randomization analysis. Front Immunol 15, 1412032 (2024).

42. Mu, Z. et al. Impact of disease-associated chromatin accessibility QTLs across immune cell types and contexts. Cell Genomics 6, 101061 (2026).

43. Mostafavi, H., Spence, J. P., Naqvi, S. & Pritchard, J. K. Systematic differences in discovery of genetic effects on gene expression and complex traits. Nat Genet 55, 1866–1875 (2023).

44. Umans, B. D., Battle, A. & Gilad, Y. Where Are the Disease-Associated eQTLs? Trends in Genetics 37, 109–124 (2021).

45. Kichaev, G. et al. Integrating Functional Data to Prioritize Causal Variants in Statistical Fine-Mapping Studies. PLOS Genetics 10, e1004722 (2014).

46. Mahajan, A. et al. Fine-mapping type 2 diabetes loci to single-variant resolution using high-density imputation and islet-specific epigenome maps. Nat Genet 50, 1505–1513 (2018).

47. Gazal, S. et al. Functional architecture of low-frequency variants highlights strength of negative selection across coding and non-coding annotations. Nat Genet 50, 1600–1607 (2018).

48. Avsec, Ž., et al. Advancing regulatory variant effect prediction with AlphaGenome. Nature 649, 1206–1218 (2026).

49. Linder, J., Srivastava, D., Yuan, H., Agarwal, V. & Kelley, D. R. Predicting RNA-seq coverage from DNA sequence as a unifying model of gene regulation. Nat Genet 57, 949–961 (2025).

50. Linder, J., Yuan, H. & Kelley, D. R. Predicting cell type-specific coverage profiles from DNA sequence. 2025.06.10.658961 Preprint at 10.1101/2025.06.10.658961 (2025).

51. Morris, J. A. et al. Discovery of target genes and pathways at GWAS loci by pooled single-cell CRISPR screens. Science 380, eadh7699 (2023).

52. Ghatan, S. et al. CRISPRi perturbation screens and eQTLs provide complementary and distinct insights into GWAS target genes. 2025.05.05.651929 Preprint at 10.1101/2025.05.05.651929 (2025).

53. Avsec, Ž., et al. Effective gene expression prediction from sequence by integrating long-range interactions. Nat Methods 18, 1196–1203 (2021).

54. Benegas, G., Albors, C., Aw, A. J., Ye, C. & Song, Y. S. A DNA language model based on multispecies alignment predicts the effects of genome-wide variants. Nat Biotechnol 43, 1960–1965 (2025).

55. Pampari, A. et al. ChromBPNet: bias factorized, base-resolution deep learning models of chromatin accessibility reveal cis-regulatory sequence syntax, transcription factor footprints and regulatory variants. bioRxiv 2024.12.25.630221 (2025) doi:10.1101/2024.12.25.630221.

56. Weiner, R. J., Lakhani, C., Knowles, D. A. & Gürsoy, G. LDmat: efficiently queryable compression of linkage disequilibrium matrices. Bioinformatics 39, btad092 (2023).

57. Bercovich, U., Zabad, S. & Gravel, S. LD Matrix Approximations for Scalable Analysis of High-dimensional Genetic Data. 2025.09.16.676478 Preprint at 10.1101/2025.09.16.676478 (2025).

58. Sanctis, G. E. de et al. SAFE-LD: A novel method for the estimation of linkage disequilibrium from summary statistics. 2025.09.29.679154 Preprint at 10.1101/2025.09.29.679154 (2025).

59. Carss, K. et al. Whole-genome sequencing of 490,640 UK Biobank participants. Nature 645, 692–701 (2025).

60. Bick, A. G. et al. Genomic data in the All of Us Research Program. Nature 627, 340–346 (2024).

61. Zaitlen, N., Paşaniuc, B., Gur, T., Ziv, E. & Halperin, E. Leveraging Genetic Variability across Populations for the Identification of Causal Variants. The American Journal of Human Genetics 86, 23–33 (2010).

62. Carss, K. et al. Whole-genome sequencing of 490,640 UK Biobank participants. Nature 645, 692–701 (2025).

63. Gao, B. & Zhou, X. MESuSiE enables scalable and powerful multi-ancestry fine-mapping of causal variants in genome-wide association studies. Nat Genet 56, 170–179 (2024).

64. Rossen, J. et al. MultiSuSiE improves multi-ancestry fine-mapping in All of Us whole-genome sequencing data. Nat Genet 58, 67–76 (2026).

65. Yuan, K. et al. Fine-mapping across diverse ancestries drives the discovery of putative causal variants underlying human complex traits and diseases. Nat Genet 56, 1841–1850 (2024).

66. Pickrell, J. K. Joint Analysis of Functional Genomic Data and Genome-wide Association Studies of 18 Human Traits. The American Journal of Human Genetics 94, 559–573 (2014).

67. Weissbrod, O. et al. Functionally informed fine-mapping and polygenic localization of complex trait heritability. Nat Genet 52, 1355–1363 (2020).

68. Srivastava, D. et al. Borzoi-informed fine mapping improves causal variant prioritization in complex trait GWAS. 2025.07.09.663936 Preprint at 10.1101/2025.07.09.663936 (2025).

69. Engreitz, J. M. et al. Deciphering the impact of genomic variation on function. Nature 633, 47–57 (2024).

70. Deng, C. et al. Massively parallel characterization of regulatory elements in the developing human cortex. Science 384, eadh0559 (2024).

71. Green, N. F. O. et al. CRISPRi screening in cultured human astrocytes uncovers distal enhancers controlling genes dysregulated in Alzheimer’s disease. Nat Neurosci 29, 703–716 (2026).

72. Morris, J. A. et al. Discovery of target genes and pathways at GWAS loci by pooled single-cell CRISPR screens. Science 380, eadh7699 (2023).

73. Nagai, M., Murphy, A. E., Rizzo, K. & Koo, P. K. Toward Interpretable and Generalizable AI in Regulatory Genomics. Preprint at 10.48550/arXiv.2602.01230 (2026).

74. Morrison, J. GWASBrewer: An R Package for Simulating Realistic GWAS Summary Statistics. Genet Epidemiol 49, e22594 (2025).

75. Benegas, G., Eraslan, G. & Song, Y. S. Benchmarking DNA Sequence Models for Causal Regulatory Variant Prediction in Human Genetics. bioRxiv 2025.02.11.637758 (2025) doi:10.1101/2025.02.11.637758.

76. Saevarsdottir, S. et al. FLT3 stop mutation increases FLT3 ligand level and risk of autoimmune thyroid disease. Nature 584, 619–623 (2020).

