## Supplementary Figures and Tables for "Current challenges in GWAS integration and fine-mapping for variant interpretation"

**Supplementary materials**

**Supplementary Figure 1.** Precision-recall of different deep learning variant effect predictors.

**Supplementary Figure 2.** Genome-wide significant variants across blood trait GWAS loci.

**Supplementary Table 1.** Full set of 123 autoimmune disease GWAS.

**Supplementary Table 2.** Number of genotyping arrays used in each GWAS.

**Supplementary Table 3.** Largest GWAS for each of the 35 distinct autoimmune diseases.

**Supplementary Table 4.** Colocalization of GWAS signal across 16 autoimmune GWAS.

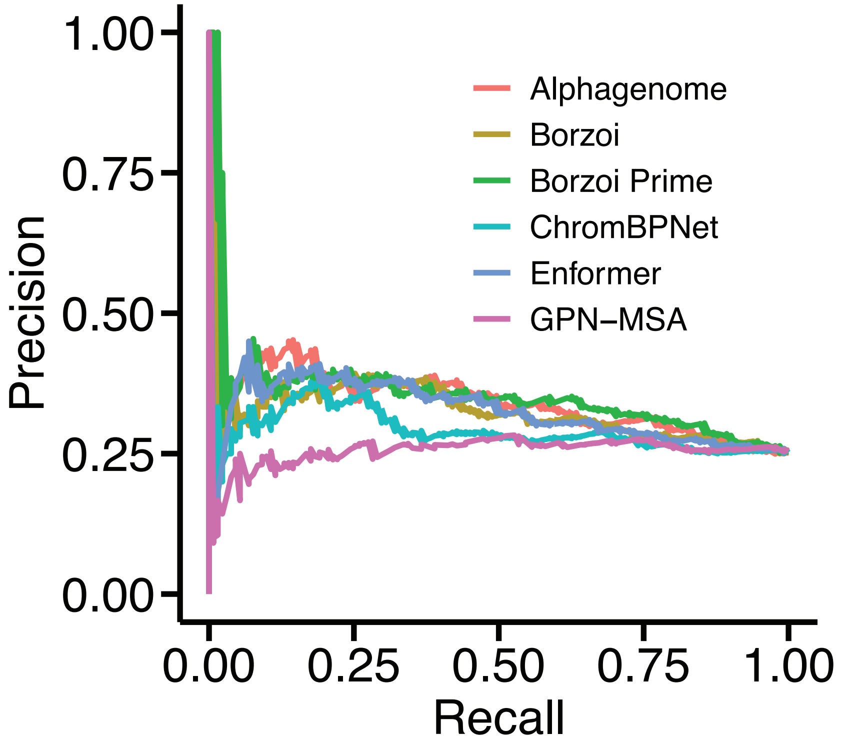


**Supplementary Fig. 1 | Precision-recall of different deep learning variant effect predictors**

We applied six different variant effect predictor tools to distinguish between the STING-positive and STING-negative variants identified by Morris et al.[1] and observed Borzoi Prime and AlphaGenome yielded the largest AUPRCs.

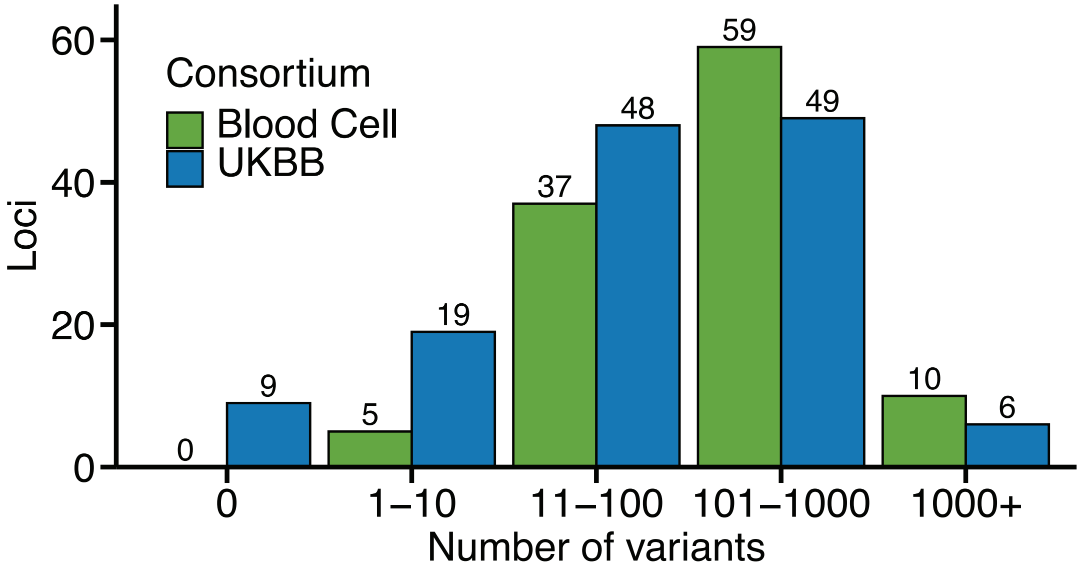


**Supplementary Fig. 2 | Genome-wide significant variants across blood trait GWAS loci**

From the Morris *et al.* perturbation study[1], we focused on the GWAS loci containing a STING-positive variant and extracted the genome-wide significant variants in the summary statistics within 500 kb from the STING-positive variant. For our scoring analyses in *Figure 5*, we only chose GWAS loci with at least 50 genome-wide significant variants so that the task of nominating the STING-positive variant would not be trivial.

**Supplementary Table 1 | Full set of 123 autoimmune disease GWAS**

| i | GWAS ID | PubMed ID | Trait abbreviation | GWAS ancestry |
|---|---|---|---|---|
| 1 | 2008_18204098_SLE_EUR | 18204098 | SLE | EUR |
| 2 | 2010_20190752_CEL_EUR | 20190752 | CEL | EUR |
| 3 | 2010_20453842_RA_EUR | 20453842 | RA | EUR |
| 4 | 2011_21833088_MS_EUR | 21833088 | MS | EUR |
| 5 | 2012_23143594_PSO_EUR | 23143594 | PSO | EUR |
| 6 | 2012_23143596_RA_EUR | 23143596 | RA | EUR |
| 7 | 2013_23603761_JIA_EUR | 23603761 | JIA | EUR |
| 8 | 2013_23749187_AS_EUR-EAS | 23749187 | AS | EUR-EAS |
| 9 | 2013_24076602_MS_EUR | 24076602 | MS | EUR |
| 10 | 2013_24390342_RA_EUR | 24390342 | RA | EUR |
| 11 | 2015_25751624_T1D_EUR | 25751624 | T1D | EUR |
| 12 | 2015_26192919_IBD_EUR | 26192919 | IBD | EUR |
| 13 | 2015_26394269_PBC_EUR | 26394269 | PBC | EUR |
| 14 | 2015_26482879_AD_EUR-EAS-AFR | 26482879 | AD | EUR-EAS-AFR |
| 15 | 2015_26502338_SLE_EUR | 26502338 | SLE | EUR |
| 16 | 2016_27386562_MS_EUR | 27386562 | MS | EUR |
| 17 | 2016_27723757_VIT_EUR | 27723757 | VIT | EUR |
| 18 | 2016_27723758_SIGD_EUR | 27723758 | SIGD | EUR |
| 19 | 2016_27992413_PSC_EUR | 27992413 | PSC | EUR |
| 20 | 2017_28067908_IBD_EUR | 28067908 | IBD | EUR |
| 21 | 2017_29083406_AT_EUR | 29083406 | AT | EUR |
| 22 | 2018_29769526_NO_EUR | 29769526 | NO | EUR |
| 23 | 2018_30254083_LADA_EUR | 30254083 | LADA | EUR |
| 24 | 2018_30343897_MS_EUR | 30343897 | MS | EUR |
| 25 | 2019_30946743_AS_MEA | 30946743 | AS | MEA |
| 26 | 2019_31604244_MS_EUR | 31604244 | MS | EUR |
| 27 | 2019_31719529_EGPA_EUR | 31719529 | EGPA | EUR |
| 28 | 2020_31959851_AST_EUR | 31959851 | AST | EUR |
| 29 | 2020_32005708_T1D_EUR | 32005708 | T1D | EUR |
| 30 | 2020_32231244EAS_MN_EAS | 32231244 | MN | EAS |
| 31 | 2020_32231244_MN_EUR-EAS | 32231244 | MN | EUR-EAS |

| | | | | |
|---|---|---|---|---|
| 32 | 2020_32296059_AST_EUR | 32296059 | AST | EUR |
| 33 | 2020_32514122_AD_EAS | 32514122 | AD | EAS |
| 34 | 2020_32514122_GD_EAS | 32514122 | GD | EAS |
| 35 | 2020_32514122_RA_EAS | 32514122 | RA | EAS |
| 36 | 2020_32581359_AITD_EUR | 32581359 | AITD | EUR |
| 37 | 2020_32888493_baso-count_EUR | 32888493 | baso-count | EUR |
| 38 | 2020_32888493_eos-count_EUR | 32888493 | eos-count | EUR |
| 39 | 2020_32888493_lymph-count_EUR | 32888493 | lymph-count | EUR |
| 40 | 2020_32888493_mono-count_EUR | 32888493 | mono-count | EUR |
| 41 | 2020_32888493_myeloid-count_EUR | 32888493 | myeloid-count | EUR |
| 42 | 2020_32888493_neutro-count_EUR | 32888493 | neutro-count | EUR |
| 43 | 2020_32888494_baso-count_EUR | 32888494 | baso-count | EUR |
| 44 | 2020_32888494_baso-pct_EUR | 32888494 | baso-pct | EUR |
| 45 | 2020_32888494_eos-count_EUR | 32888494 | eos-count | EUR |
| 46 | 2020_32888494_eos-pct_EUR | 32888494 | eos-pct | EUR |
| 47 | 2020_32888494_lymph-count_EUR | 32888494 | lymph-count | EUR |
| 48 | 2020_32888494_lymph-pct_EUR | 32888494 | lymph-pct | EUR |
| 49 | 2020_32888494_mono-count_EUR | 32888494 | mono-count | EUR |
| 50 | 2020_32888494_mono-pct_EUR | 32888494 | mono-pct | EUR |
| 51 | 2020_32888494_neutro-count_EUR | 32888494 | neutro-count | EUR |
| 52 | 2020_32888494_neutro-pct_EUR | 32888494 | neutro-pct | EUR |
| 53 | 2020_33067605_CCL20_EUR | 33067605 | CCL20 | EUR |
| 54 | 2020_33067605_CCL4_EUR | 33067605 | CCL4 | EUR |
| 55 | 2020_33067605_CD40_EUR | 33067605 | CD40 | EUR |
| 56 | 2020_33067605_CXCL1_EUR | 33067605 | CXCL1 | EUR |
| 57 | 2020_33067605_IL18_EUR | 33067605 | IL18 | EUR |
| 58 | 2020_33067605_IL1_EUR | 33067605 | IL1 | EUR |
| 59 | 2020_33067605_IL27_EUR | 33067605 | IL27 | EUR |
| 60 | 2020_33067605_IL6_EUR | 33067605 | IL6 | EUR |
| 61 | 2020_33067605_IL8_EUR | 33067605 | IL8 | EUR |
| 62 | 2020_33067605_MCSF_EUR | 33067605 | MCSF | EUR |
| 63 | 2020_33067605_TNFR1_EUR | 33067605 | TNFR1 | EUR |
| 64 | 2020_33067605_TNFR2_EUR | 33067605 | TNFR2 | EUR |
| 65 | 2020_33067605_TNF_EUR | 33067605 | TNF | EUR |

| | | | | |
|---|---|---|---|---|
| 66 | 2020_33067605_TRAILR2_EUR | 33067605 | TRAILR2 | EUR |
| 67 | 2020_33067605_mono-count_EUR | 33067605 | mono-count | EUR |
| 68 | 2020_33067605_proIL16_EUR | 33067605 | proIL16 | EUR |
| 69 | 2020_33106285_JIA_EUR | 33106285 | JIA | EUR |
| 70 | 2020_33310728_RA_EUR-EAS | 33310728 | RA | EUR-EAS |
| 71 | 2021_33493351_SLE_EAS | 33493351 | SLE | EAS |
| 72 | 2021_33536424EAS_SLE_EAS | 33536424 | SLE | EAS |
| 73 | 2021_33536424_SLE_EUR-EAS | 33536424 | SLE | EUR-EAS |
| 74 | 2021_33574239_ADD_EUR | 33574239 | ADD | EUR |
| 75 | 2021_34012112_T1D_EUR | 34012112 | T1D | EUR |
| 76 | 2021_34127860_T1D_EUR | 34127860 | T1D | EUR |
| 77 | 2021_34594039EAS_AD_EAS | 34594039 | AD | EAS |
| 78 | 2021_34594039EAS_AST_EAS | 34594039 | AST | EAS |
| 79 | 2021_34594039EAS_CS_EAS | 34594039 | CS | EAS |
| 80 | 2021_34594039EAS_GD_EAS | 34594039 | GD | EAS |
| 81 | 2021_34594039EAS_RA_EAS | 34594039 | RA | EAS |
| 82 | 2021_34594039EAS_T1D_EAS | 34594039 | T1D | EAS |
| 83 | 2021_34594039_AD_EUR-EAS | 34594039 | AD | EUR-EAS |
| 84 | 2021_34594039_AST_EUR-EAS | 34594039 | AST | EUR-EAS |
| 85 | 2021_34594039_CS_EUR-EAS | 34594039 | CS | EUR-EAS |
| 86 | 2021_34594039_GD_EUR-EAS | 34594039 | GD | EUR-EAS |
| 87 | 2021_34594039_HT_EUR-EAS | 34594039 | HT | EUR-EAS |
| 88 | 2021_34594039_IBD_EUR-EAS | 34594039 | IBD | EUR-EAS |
| 89 | 2021_34594039_IGN_EUR-EAS | 34594039 | IGN | EUR-EAS |
| 90 | 2021_34594039_PDAST_EUR-EAS | 34594039 | PDAST | EUR-EAS |
| 91 | 2021_34594039_PV_EUR-EAS | 34594039 | PV | EUR-EAS |
| 92 | 2021_34594039_RA_EUR-EAS | 34594039 | RA | EUR-EAS |
| 93 | 2021_34594039_SD_EUR-EAS | 34594039 | SD | EUR-EAS |
| 94 | 2021_34594039_T1D_EUR-EAS | 34594039 | T1D | EUR-EAS |
| 95 | 2022_35074870_MG_EUR | 35074870 | MG | EUR |
| 96 | 2022_35470158_RA_EUR | 35470158 | RA | EUR |
| 97 | 2022_35507331_PSO_EUR | 35507331 | PSO | EUR |
| 98 | 2022_35896530_SD_EUR | 35896530 | SD | EUR |
| 99 | 2022_36333501EAS_RA_EAS | 36333501 | RA | EAS |

| | | | | |
|---|---|---|---|---|
| 100 | 2022_36333501_RA_EUR | 36333501 | RA | EUR |
| 101 | 2023_36750564_SLE_EUR-EAS-AMR | 36750564 | SLE | EUR-EAS-AMR |
| 102 | 2023_36828809_PSC_EUR | 36828809 | PSC | EUR |
| 103 | 2023_37156999EAS_CD_EAS | 37156999 | CD | EAS |
| 104 | 2023_37156999EAS_IBD_EAS | 37156999 | IBD | EAS |
| 105 | 2023_37156999EAS_UC_EAS | 37156999 | UC | EAS |
| 106 | 2023_37156999_CD_EUR-EAS | 37156999 | CD | EUR-EAS |
| 107 | 2023_37156999_IBD_EUR-EAS | 37156999 | IBD | EUR-EAS |
| 108 | 2023_37156999_UC_EUR-EAS | 37156999 | UC | EUR-EAS |
| 109 | 2023_37337107_IGN_EUR-EAS | 37337107 | IGN | EUR-EAS |
| 110 | 2023_37752970_ALO_EAS | 37752970 | ALO | EAS |
| 111 | 2024_38296975_SS_EAS | 38296975 | SS | EAS |
| 112 | 2024_38982041_AITD_EUR | 38982041 | AITD | EUR |
| 113 | 2024_39537604_MG_EUR | 39537604 | MG | EUR |
| 114 | 2024_PanUKBB_AST_EUR | 40968291 | AST | EUR |
| 115 | 2024_PanUKBB_AS_EUR | 40968291 | AS | EUR |
| 116 | 2024_PanUKBB_CD_EUR | 40968291 | CD | EUR |
| 117 | 2024_PanUKBB_ECZ_EUR | 40968291 | ECZ | EUR |
| 118 | 2024_PanUKBB_IBD_EUR | 40968291 | IBD | EUR |
| 119 | 2024_PanUKBB_PSO_EUR | 40968291 | PSO | EUR |
| 120 | 2024_PanUKBB_RA_EUR | 40968291 | RA | EUR |
| 121 | 2024_PanUKBB_SLE_EUR | 40968291 | SLE | EUR |
| 122 | 2024_PanUKBB_T1D_EUR | 40968291 | T1D | EUR |
| 123 | 2024_PanUKBB_UC_EUR | 40968291 | UC | EUR |

**Supplementary Table 2 | Number of genotyping arrays used in each GWAS**

| i | GWAS ID | Genome | Number of genotyping arrays |
|---|---|---|---|
| 1 | 2023_36750564_SLE_EUR-EAS-AMR | GRCh37 | 3+ |
| 2 | 2022_36333501_RA_EUR | GRCh37 | 3+ |
| 3 | 2015_26192919_IBD_EUR | GRCh37 | 3+ |
| 4 | 2013_24076602_MS_EUR | GRCh37 | 1 |
| 5 | 2020_33310728_RA_EUR-EAS | GRCh37 | 3+ |
| 6 | 2011_21833088_MS_EUR | GRCh37 | 2 |
| 7 | 2016_27723757_VIT_EUR | GRCh37 | 1 |
| 8 | 2015_25751624_T1D_EUR | GRCh37 | 1 |
| 9 | 2015_26502338_SLE_EUR | GRCh37 | 3+ |
| 10 | 2023_37337107_IGN_EUR-EAS | GRCh38 | 3+ |
| 11 | 2012_23143594_PSO_EUR | GRCh37 | 1 |
| 12 | 2010_20190752_CEL_EUR | GRCh37 | 3+ |
| 13 | 2021_33536424_SLE_EUR-EAS | GRCh37 | 3+ |
| 14 | 2016_27992413_PSC_EUR | GRCh37 | 3+ |
| 15 | 2010_20453842_RA_EUR | GRCh37 | 3+ |
| 16 | 2016_27386562_MS_EUR | GRCh37 | 3+ |
| 17 | 2022_35507331_PSO_EUR | GRCh37 | 3+ |
| 18 | 2020_32005708_T1D_EUR | GRCh37 | 3+ |
| 19 | 2019_31719529_EGPA_EUR | GRCh38 | 3+ |
| 20 | 2019_31604244_MS_EUR | GRCh38 | 2 |
| 21 | 2017_28067908_IBD_EUR | GRCh37 | 2 |
| 22 | 2021_34127860_T1D_EUR | GRCh38 | 1 |
| 23 | 2021_34012112_T1D_EUR | GRCh38 | 3+ |
| 24 | 2022_35074870_MG_EUR | GRCh38 | 3+ |
| 25 | 2013_24390342_RA_EUR | GRCh37 | 3+ |
| 26 | 2019_30946743_AS_MEA | GRCh37 | 2 |
| 27 | 2024_38982041_AITD_EUR | GRCh38 | 3+ |
| 28 | 2022_35470158_RA_EUR | GRCh38 | 2 |
| 29 | 2023_36828809_PSC_EUR | GRCh37 | 3+ |
| 30 | 2020_31959851_AST_EUR | GRCh38 | 3+ |
| 31 | 2020_32581359_AITD_EUR | GRCh38 | 2 |
| 32 | 2020_32231244_MN_EUR-EAS | GRCh37 | 3+ |

| | | | |
|---|---|---|---|
| 33 | 2021_34594039_SD_EUR-EAS | GRCh37 | 3+ |
| 34 | 2021_34594039_GD_EUR-EAS | GRCh37 | 3+ |
| 35 | 2021_34594039_PDAST_EUR-EAS | GRCh37 | 3+ |
| 36 | 2021_34594039_AST_EUR-EAS | GRCh37 | 3+ |
| 37 | 2021_34594039_RA_EUR-EAS | GRCh37 | 3+ |
| 38 | 2021_34594039_IBD_EUR-EAS | GRCh37 | 3+ |
| 39 | 2021_34594039_AD_EUR-EAS | GRCh37 | 3+ |
| 40 | 2021_34594039_HT_EUR-EAS | GRCh37 | 3+ |
| 41 | 2021_34594039_T1D_EUR-EAS | GRCh37 | 3+ |
| 42 | 2021_34594039_PV_EUR-EAS | GRCh37 | 3+ |
| 43 | 2021_34594039_IGN_EUR-EAS | GRCh37 | 3+ |
| 44 | 2021_34594039_CS_EUR-EAS | GRCh37 | 3+ |
| 45 | 2023_37156999_IBD_EUR-EAS | GRCh38 | 3+ |
| 46 | 2023_37156999_CD_EUR-EAS | GRCh38 | 3+ |
| 47 | 2023_37156999_UC_EUR-EAS | GRCh38 | 3+ |
| 48 | 2013_23749187_AS_EUR-EAS | GRCh37 | 3+ |
| 49 | 2024_39537604_MG_EUR | GRCh37 | 3+ |
| 50 | 2018_29769526_NO_EUR | GRCh37 | 3+ |
| 51 | 2013_23603761_JIA_EUR | GRCh37 | 1 |
| 52 | 2020_33106285_JIA_EUR | GRCh37 | 3+ |
| 53 | 2018_30254083_LADA_EUR | GRCh37 | 3+ |
| 54 | 2015_26482879_AD_EUR-EAS-AFR | GRCh37 | 3+ |
| 55 | 2017_29083406_AT_EUR | GRCh37 | 3+ |
| 56 | 2020_32296059_AST_EUR | GRCh37 | 3+ |
| 57 | 2016_27723758_SIGD_EUR | GRCh37 | 3+ |
| 58 | 2012_23143596_RA_EUR | GRCh37 | 1 |
| 59 | 2018_30343897_MS_EUR | GRCh37 | 2 |
| 60 | 2021_33493351_SLE_EAS | GRCh37 | 1 |
| 61 | 2008_18204098_SLE_EUR | GRCh36 | 2 |
| 62 | 2015_26394269_PBC_EUR | GRCh37 | 2 |
| 63 | 2023_37752970_ALO_EAS | GRCh38 | 1 |
| 64 | 2024_38296975_SS_EAS | GRCh37 | 3+ |
| 65 | 2022_35896530_SD_EUR | GRCh38 | 1 |
| 66 | 2020_32514122_GD_EAS | GRCh37 | 2 |

| | | | |
|---|---|---|---|
| 67 | 2020_32514122_RA_EAS | GRCh37 | 2 |
| 68 | 2020_32514122_AD_EAS | GRCh37 | 2 |
| 69 | 2024_PanUKBB_SLE_EUR | GRCh37 | 3+ |
| 70 | 2024_PanUKBB_AST_EUR | GRCh37 | 3+ |
| 71 | 2024_PanUKBB_T1D_EUR | GRCh37 | 3+ |
| 72 | 2024_PanUKBB_AS_EUR | GRCh37 | 3+ |
| 73 | 2024_PanUKBB_ECZ_EUR | GRCh37 | 3+ |
| 74 | 2024_PanUKBB_PSO_EUR | GRCh37 | 3+ |
| 75 | 2024_PanUKBB_IBD_EUR | GRCh37 | 3+ |
| 76 | 2024_PanUKBB_CD_EUR | GRCh37 | 3+ |
| 77 | 2024_PanUKBB_UC_EUR | GRCh37 | 3+ |
| 78 | 2024_PanUKBB_RA_EUR | GRCh37 | 3+ |
| 79 | 2020_32231244EAS_MN_EAS | GRCh37 | 1 |
| 80 | 2021_33536424EAS_SLE_EAS | GRCh37 | 1 |
| 81 | 2021_34594039EAS_GD_EAS | GRCh37 | 2 |
| 82 | 2021_34594039EAS_AST_EAS | GRCh37 | 2 |
| 83 | 2021_34594039EAS_RA_EAS | GRCh37 | 2 |
| 84 | 2021_34594039EAS_AD_EAS | GRCh37 | 2 |
| 85 | 2021_34594039EAS_T1D_EAS | GRCh37 | 2 |
| 86 | 2021_34594039EAS_CS_EAS | GRCh37 | 2 |
| 87 | 2023_37156999EAS_IBD_EAS | GRCh38 | 1 |
| 88 | 2023_37156999EAS_CD_EAS | GRCh38 | 1 |
| 89 | 2023_37156999EAS_UC_EAS | GRCh38 | 1 |
| 90 | 2022_36333501EAS_RA_EAS | GRCh37 | 1 |
| 91 | 2021_33574239_ADD_EUR | GRCh37 | 3+ |
| 92 | 2020_33067605_IL8_EUR | GRCh37 | 3+ |
| 93 | 2020_33067605_CD40_EUR | GRCh37 | 3+ |
| 94 | 2020_33067605_IL1_EUR | GRCh37 | 3+ |
| 95 | 2020_33067605_IL6_EUR | GRCh37 | 3+ |
| 96 | 2020_33067605_mono-count_EUR | GRCh37 | 3+ |
| 97 | 2020_33067605_TNF_EUR | GRCh37 | 3+ |
| 98 | 2020_33067605_TNFR1_EUR | GRCh37 | 3+ |
| 99 | 2020_33067605_IL27_EUR | GRCh37 | 3+ |
| 100 | 2020_33067605_MCSF_EUR | GRCh37 | 3+ |

| | | | |
|---|---|---|---|
| 101 | 2020_33067605_CXCL1_EUR | GRCh37 | 3+ |
| 102 | 2020_33067605_TRAILR2_EUR | GRCh37 | 3+ |
| 103 | 2020_33067605_IL18_EUR | GRCh37 | 3+ |
| 104 | 2020_33067605_TNFR2_EUR | GRCh37 | 3+ |
| 105 | 2020_33067605_proIL16_EUR | GRCh37 | 3+ |
| 106 | 2020_33067605_CCL4_EUR | GRCh37 | 3+ |
| 107 | 2020_33067605_CCL20_EUR | GRCh37 | 3+ |
| 108 | 2020_32888493_baso-count_EUR | GRCh37 | 3+ |
| 109 | 2020_32888493_eos-count_EUR | GRCh37 | 3+ |
| 110 | 2020_32888493_lymph-count_EUR | GRCh37 | 3+ |
| 111 | 2020_32888493_mono-count_EUR | GRCh37 | 3+ |
| 112 | 2020_32888493_neutro-count_EUR | GRCh37 | 3+ |
| 113 | 2020_32888493_myeloid-count_EUR | GRCh37 | 3+ |
| 114 | 2020_32888494_baso-count_EUR | GRCh37 | 3+ |
| 115 | 2020_32888494_baso-pct_EUR | GRCh37 | 3+ |
| 116 | 2020_32888494_eos-count_EUR | GRCh37 | 3+ |
| 117 | 2020_32888494_eos-pct_EUR | GRCh37 | 3+ |
| 118 | 2020_32888494_lymph-count_EUR | GRCh37 | 3+ |
| 119 | 2020_32888494_lymph-pct_EUR | GRCh37 | 3+ |
| 120 | 2020_32888494_mono-count_EUR | GRCh37 | 3+ |
| 121 | 2020_32888494_mono-pct_EUR | GRCh37 | 3+ |
| 122 | 2020_32888494_neutro-count_EUR | GRCh37 | 3+ |
| 123 | 2020_32888494_neutro-pct_EUR | GRCh37 | 3+ |

**Supplementary Table 3 | Largest GWAS for each of the 35 distinct autoimmune diseases**

| i | GWAS ID | PubMed ID | Trait abbreviation | GWAS ancestry |
|---|---|---|---|---|
| 1 | 2010_20190752_CEL_EUR | 20190752 | CEL | EUR |
| 2 | 2012_23143594_PSO_EUR | 23143594 | PSO | EUR |
| 3 | 2013_23749187_AS_EUR-EAS | 23749187 | AS | EUR-EAS |
| 4 | 2015_26394269_PBC_EUR | 26394269 | PBC | EUR |
| 5 | 2016_27723757_VIT_EUR | 27723757 | VIT | EUR |
| 6 | 2016_27723758_SIGD_EUR | 27723758 | SIGD | EUR |
| 7 | 2016_27992413_PSC_EUR | 27992413 | PSC | EUR |
| 8 | 2017_29083406_AT_EUR | 29083406 | AT | EUR |
| 9 | 2018_29769526_NO_EUR | 29769526 | NO | EUR |
| 10 | 2018_30254083_LADA_EUR | 30254083 | LADA | EUR |
| 11 | 2019_31604244_MS_EUR | 31604244 | MS | EUR |
| 12 | 2019_31719529_EGPA_EUR | 31719529 | EGPA | EUR |
| 13 | 2020_32231244_MN_EUR-EAS | 32231244 | MN | EUR-EAS |
| 14 | 2020_32296059_AST_EUR | 32296059 | AST | EUR |
| 15 | 2020_33106285_JIA_EUR | 33106285 | JIA | EUR |
| 16 | 2020_33310728_RA_EUR-EAS | 33310728 | RA | EUR-EAS |
| 17 | 2021_33574239_ADD_EUR | 33574239 | ADD | EUR |
| 18 | 2021_34012112_T1D_EUR | 34012112 | T1D | EUR |
| 19 | 2021_34594039_AD_EUR-EAS | 34594039 | AD | EUR-EAS |
| 20 | 2021_34594039_CS_EUR-EAS | 34594039 | CS | EUR-EAS |
| 21 | 2021_34594039_GD_EUR-EAS | 34594039 | GD | EUR-EAS |
| 22 | 2021_34594039_HT_EUR-EAS | 34594039 | HT | EUR-EAS |
| 23 | 2021_34594039_IGN_EUR-EAS | 34594039 | IGN | EUR-EAS |
| 24 | 2021_34594039_PDAST_EUR-EAS | 34594039 | PDAST | EUR-EAS |
| 25 | 2021_34594039_PV_EUR-EAS | 34594039 | PV | EUR-EAS |
| 26 | 2022_35896530_SD_EUR | 35896530 | SD | EUR |
| 27 | 2023_36750564_SLE_EUR-EAS-AMR | 36750564 | SLE | EUR-EAS-AMR |
| 28 | 2023_37156999_CD_EUR-EAS | 37156999 | CD | EUR-EAS |
| 29 | 2023_37156999_IBD_EUR-EAS | 37156999 | IBD | EUR-EAS |
| 30 | 2023_37156999_UC_EUR-EAS | 37156999 | UC | EUR-EAS |
| 31 | 2023_37752970_ALO_EAS | 37752970 | ALO | EAS |
| 32 | 2024_38296975_SS_EAS | 38296975 | SS | EAS |

| | | | | |
|---|---|---|---|---|
| 33 | 2024_38982041_AITD_EUR | 38982041 | AITD | EUR |
| 34 | 2024_39537604_MG_EUR | 39537604 | MG | EUR |
| 35 | 2024_PanUKBB_ECZ_EUR | PanUKBB | ECZ | EUR |

**Supplementary Table 4 | Colocalization of GWAS signal across 16 autoimmune GWAS**

| i | GWAS ID | Trait abbreviation | Contains genome-wide significant variant? |
|---|---|---|---|
| 1 | 2020_32514122_GD_EAS | GD | Yes |
| 2 | 2021_34594039EAS_GD_EAS | GD | Yes |
| 3 | 2024_38982041_AITD_EUR | AITD | Yes |
| 4 | 2021_34594039_GD_EUR-EAS | GD | Yes |
| 5 | 2020_32581359_AITD_EUR | AITD | Yes |
| 6 | 2016_27723757_VIT_EUR | VIT | Yes |
| 7 | 2019_31604244_MS_EUR | MS | No |
| 8 | 2020_32005708_T1D_EUR | T1D | No |
| 9 | 2022_36333501EAS_RA_EAS | RA | No |
| 10 | 2020_33310728_RA_EUR-EAS | RA | No |
| 11 | 2021_34594039_RA_EUR-EAS | RA | No |
| 12 | 2022_35470158_RA_EUR | RA | No |
| 13 | 2021_34012112_T1D_EUR | T1D | No |
| 14 | 2020_33106285_JIA_EUR | JIA | No |
| 15 | 2023_37156999_CD_EUR-EAS | CD | No |
| 16 | 2016_27723758_SIGD_EUR | SIGD | No |

### References

1. Morris, J. A. *et al.* Discovery of target genes and pathways at GWAS loci by pooled single-cell CRISPR screens. *Science* **380**, eadh7699 (2023).
